# A high throughput screen with a clonogenic endpoint to identify radiation modulators of cancer

**DOI:** 10.1101/2022.05.24.493331

**Authors:** Nathan P. Gomes, Barbara Frederick, Jeremy R. Jacobsen, Doug Chapnick, Tin Tin Su

**Affiliations:** SuviCa, Inc. PO Box 3131, Boulder CO, 80307-3131; Department of Molecular, Cellular and Developmental Biology, University of Colorado Boulder, CO, 80309-0347; Molecular and Cellular Oncology Program, University of Colorado Cancer Center, Aurora, CO 80045; BioLoomics, 4665 Nautilus Ct. S., Ste. 501, Boulder, CO 80301

## Abstract

Gomes, N. P., Frederick, B., Jacobsen, J. R., Chapnick. D. and Su, T. T. A high throughput screen with a clonogenic endpoint to identify radiation modulators of cancer. *Radiat. Res*.

Clonogenic assays evaluate the ability of single cells to proliferate and form colonies. This process approximates the regrowth and recurrence of tumors after treatment with radiation or chemotherapy, and thereby provides a drug discovery platform for compounds that block this process. However, because of their labor-intensive and cumbersome nature, adapting canonical clonogenic assays for high throughput screening (HTS) has been challenging. We overcame these barriers by developing an integrated system that automates cell- and liquid-handling, irradiation, dosimetry, drug administration, and incubation. Further, we developed a fluorescent live-cell based automated colony scoring methodology that identifies and counts colonies precisely based upon actual nuclei number rather than colony area, thereby eliminating errors in colony counts caused by radiation induced changes in colony morphology. We identified 13 cell lines from 7 cancer types, where radiation is a standard treatment module, that exhibit identical radiation and chemoradiation response regardless of well format and are amenable to miniaturization into small-well HTS formats. We performed pilot screens through a 1584 compound NCI Diversity Set library using two cell lines representing different cancer indications. Radiation modulators identified in the pilot screens were validated in traditional clonogenic assays, providing proof-of-concept for the screen. The integrated methodology, hereafter ‘clonogenic HTS’, exhibits excellent robustness (Z’ values >0.5) and shows high reproducibility (>95%). We propose that clonogenic HTS we developed can function as a drug discovery platform to identify compounds that inhibit tumor regrowth following radiation therapy, to identify new efficacious pair-wise combinations of known oncologic therapies, or to identify novel modulators of approved therapies.

## Introduction

More than half of US cancer patients undergo radiation therapy (RT) alone, or in combination with other oncologic therapies (*1*). Cancer recurrence following RT is common, resulting from proliferation of tumor clonogens that survived initial RT modalities (*2*). Identification of therapies that specifically inhibit clonogenic proliferation following radiation would be a significant advancement for RT-based cancer treatment, likely to reduce recurrence rates, and improve overall outcomes. Several therapeutic agents are approved for combinatorial use with RT for the treatment cancer, but none were initially identified as specific inhibitors of clonogenic growth following RT. Rather, mostly cytotoxic agents approved as discrete therapies are combined with RT empirically (*3*). It remains to be determined if direct screening for compounds that inhibit clonogenic proliferation can objectively identify novel and more potent radiation modulators. Advancements in cheminformatics, chemical synthesis, and improved ability to extract natural product compounds provide abundant and diverse chemical libraries from which to search. To systematically screen for inhibitors of clonogenic growth following RT, canonical clonogenic assays would have to be adapted to high-throughput screen (HTS) platforms, hereafter ‘clonogenic HTS’.

The tradiational clonogenic assay is the only methodology that measures growth delay and total clonogenic death, whether it be it by mitotic catastrophe, apoptosis, necrosis, terminal differentiation, reproductive death, or other modes of cellular death (*4*). It is the assay of choice for monitoring celluar growth in response to RT (*5*). Relative to simpler MTT-type HTS assays, clongenic HTS assays, in principle, would be better suited for identifing novel radiation modulators, as well as more accurately testing known-drug/RT combinations. Resulting hits from this methodology should improve success in subsequent pre-clinical studies and accelerate translation into clinically relevant therapeutics. However, traditional clonogenic assays are extremely labor-, resource- and time-intensive. The traditional requirement to laboriously manually count a minimium of 50 well-seperated colonies with greater than 50 cells per colony, for each treatment condidtion has traditionally necessitated use of lager well formats, making the assay imcompatable with known HTS methodolgies. To our knowledge no HTS system exists that follows the traditional clonogenic methodology, despite description of the original assay 6 decades ago (*6*).

Several recent papers describe methodologies that approximate clonongic assays in HTS formats by miniaturizing to small well formats and utilizing automated image analysis to score fixed and stained colonies (*7-11*). The closest approximation of a clonogenic HTS among this body of work is a ‘High Content’ screen of Non Small Cell Lung Cancer cells in 96-well plates (*10*). Cells were seeded at non-clonogenic desities that yield somewhat similar survival and radiation response as in traditional 6-well clonogenic assays. Cells were treated with radiation in the absence or presence of serially diluted drugs, and “colonies” were counted after 4 days. Assay processing was as for the conventional methodology: “colonies” were fixed, stained, destained, and scored by automatated microscopy aided image analysis. A subset of hits from the screen of 146 compounds were validated in conventional clonogenic assays, and xenograft models. While this work indicates that clongenic-like HTS screens are possible, the assay does not have time/growth periods representative of true clonogenic endpoints, and the protocol is as labor-intensive as conventional clonogenic assays, including sample processing, and the necessity for multiple drug and radiation doses. The data analysis methodology was identical to that of the conventional assay, and not amenable to large datasets produced from an HTS. In a related study, short term (5-day) survival at a single radiation dose and its modification by 18 drugs was assessed in 32 cancer cell lines (*12*). Survival was found to correlate with dose-modifying factors in traditional clonogenic assays, suggesting that a simpler, shorter-term assay in a high-throughput format may be used as the primary screen to identify potential radiation modulators, later to be validated with traditional clonogenic assays. We again note that this technology does not represent an HTS with a true clonogenic endpoint. A more recent paper reports a High Content Screen (HCS) based on the growth of cancer cells in 2D and in 3D (spheroids) assays (*8*). Colony growth of SW620 colorectal cancer cells was monitored following serial seeding dilutions to achieve 1-100 colonies per well, in the presence or absence of drug. This method allows for the testing of a small number of molecules at different plating densities, but which densities in 96-well exhibits similar growth characteristics and drug-response as cells seeded as for traditional clonogenic assays was not addressed.

The goal of HTS is to use automation to rapidly screen large numbers of compounds while reducing time, reagents, and sample manipulation compared to conventional assays. Further, automation inherently increases experimental consistency by reproducibly implementing standardized protocols. Assay formats should be amenable to handling many-fold more compounds per unit time than conventional assays. A survey of the literature indicates that most components of conventional clonogenic assays have been successfully modified for use in HTS formats, and key technological components for the assay are commercially available. However, a true clonogenic HTS remains absent; its development required the functional combination of these known methodologies/technolgoies, development of missing components, and integration into a discrete HTS sytem, where scored hits validate in traditional assays. We have accomplished these tasks and describe here an integrated high-throughput system with a clonogenic endpoint. Our system integrates and automates cell- and liquid-handling, irradiation, dosimetry, drug-addition, and incubation. We report a fluorescent live-cell based automated colony scoring methodology that identifies and counts colonies based upon precise nuclei counting rather than colony area. We report the identification of 13 cell lines, from 7 cancer types where radiation is a standard treatment module, that exhibit equivalent radiation and chemoradiation response regardless of well format and are amenable to miniaturization into small-well HTS formats. We performed pilot screens in two cell lines representing two different cancer indications with a 1584 compound library, identifying radiation modulators that validated in traditional clonogenic assays. Finally, our clonogenic HTS methodology exhibits excellent robustness (Z’>0.5) and high reproducibility (>95%).

## Results

### Identification of cell lines amenable to miniaturized clonogenic growth

Clonogenic assays are traditionally performed in 6-well plates or larger vessels, allowing for growth of single cells into colonies without significant influence from neighboring clones. The large growth area allows for well separated single cell seeding, while also achieving a minimum of 50 colonies per well, as is recommended in the literature (*13*). High throughput screens require assays to be performed in smaller well formats to allow for many drugs to be screened simultaneously. To develop a true clonogenic HTS, we set out to identify cell lines that: 1) could be grown in smaller well formats, yielding well separated colonies that number >50/well over the traditional clonogenic growth period of ∼10 days and 2) exhibit equivalent radiation response between smaller well and traditional 6-well clonogenic formats. We focused on 7 cancer indications where radiation is a main treatment modality: HNSCC, NSCLC, metastatic melanoma, glioma, soft tissue sarcoma, esophageal, and rectal cancers. Not all cancer cell lines are expected to meet all two criteria. As shown in Figure 1A, four different cell lines were seeded at clonogenic density in 6-well plates and allowed to grow for 10 days. Each colony contains >50 cells yet A2068 melanoma cells form large diffuse colonies while SW837 rectal cancer cells form discrete compact colonies. An extensive literature search for clonogenic images was performed to identified cell lines that display similar colony morphology to SW837 and were commercially available with limited use restrictions. We obtained 2 to 5 cell lines for each of the 7 cancer indications, seeded them at equivalent growth densities in 6-, 24-and 96-well plates, and grew then over a period of 7-10 days to confirm the published colony morphology (Figure 1B). Of the four representative cell lines shown, the first two formed into well-separated colonies of >50 per well in 24-well plates, while the final two did so in both 24- and 96-well plates. In general, about half of the cell lines tested met this criterion in 24-well or smaller formats.

**Figure 1.**
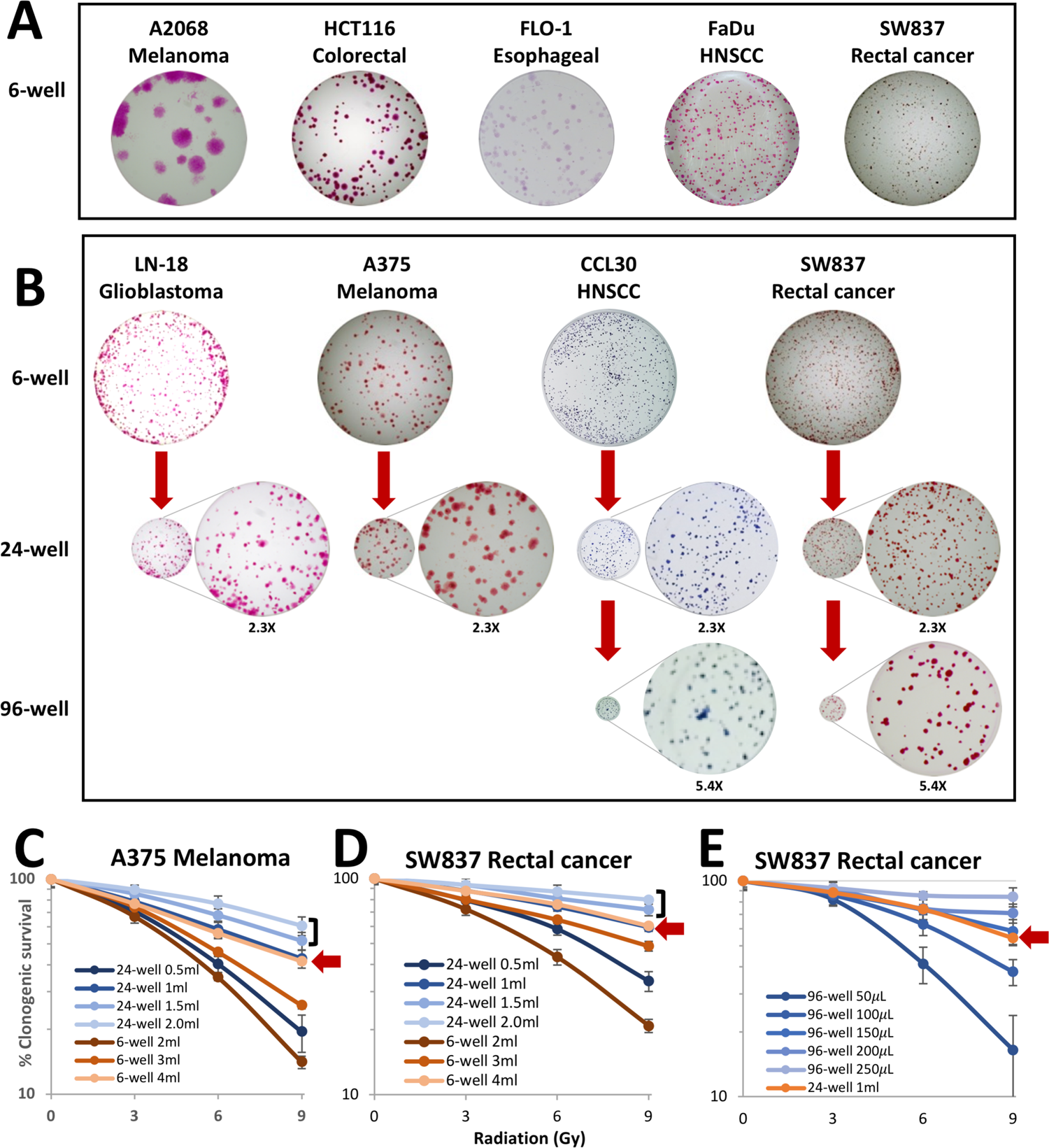
Identification of cell lines that are amenable to clonogenic growth in a small well format. (A) Five cell lines that show the variation in colony size. Each was grown for 10 days to reach colonies of 50 or more cells. (B) Growth in 6, 24 and 96 well formats. 24 well images are shown at 2.3x magnification to display the same final area as 6 well. 96 well growth is magnified 5.4x times to show equivalent area as the 6 well format. (C-E) A375 metastatic melanoma and SW837 rectal adenocarcinoma cells were seeded at the same relative density (cells/surface area) but in different volumes per well in 6-, 24-or 96-well plates. Cells were irradiated with 3, 6 or 9 Gy 24 hours after seeding. Clonogenic survival was quantified 10 days (SW837) or 7 days (A375) post radiation.

We next compared clonogenic radiation response between different well formats, using 3 doses of radiation as was done for validation of other high-content clonogenic assay formats (*10*). Figure 1C-E shows representative data for two cell lines. Cells were seeded at identical density (number of cells per growth area in the well) in the different well formats. In the traditional 6-well clonogenic assay, cells are typically grown in 4-5 ml of media per well. This volume is equivalent to 1.7-2.2 ml per well in 24-well plates and 0.7-0.9 ml per well in 96-well plates. At these equivalent media volumes, cells exhibit greater radiation resistance consistently in smaller well formats compared to 6-well plates. For example, compare ‘6-well 4ml’ (red arrow) to ‘24-well 1.5 ml’ and ‘24-well 2 ml’ (bracket) in Figure 1C-D. Systematic analysis shows that radiation survival is exquisitely sensitive to culture volume regardless of well format, with smaller relative volumes correlated with increased radiation sensitivity. We note that this is not an artifact of smaller well formats, as increased radiation sensitivity in smaller culture volumes is also seen in the traditional 6-well plates for all cell lines examined (Figure 1C, compare 6-well 2, 3, and 4 ml). One possible explanation is RT-induced production of secreted growth inhibitory factors that would be, by definition, more concentrated in smaller volumes. Importantly, as this parameter is typically not reported, variable culture volumes could partially explain lab-to-lab clonogenic variability widely observed in literature (*14*).

Regardless, consistent variation of radiation survival relative to culture volume allowed for the identification of conditions where survival in smaller well formats are equivalent to that of traditional 6-well formats. Specifically, cells grown in 4 ml in 6-well plates, 1 ml in 24-well plates and 150 μl in 96-well plates produced equivalent radiation survival curves, with ‘equivalent’ defined as average survival from technical replicates that fall within one standard deviation at each radiation dose tested (red arrows in Figure 1C-E). Similar results were obtained for all cell lines tested (thirteen total), suggesting that this phenomenon is specific to well format and seeding volume but independent of the cell line used. More specifically, nine of thirteen cell lines showed equivalent radiation survival in 6-well and 24-well formats when culture volumes were controlled while the remaining four showed equivalent radiation survival in 6-well, 24-well and 96-well formats when culture volumes were controlled (Table 1 and Supplemental Figure 1). We identified two cell lines for each of the seven cancer indications of interest, except for glioma where only one of more than ten cell lines we examined was amenable to miniaturization.

**Table 1.**
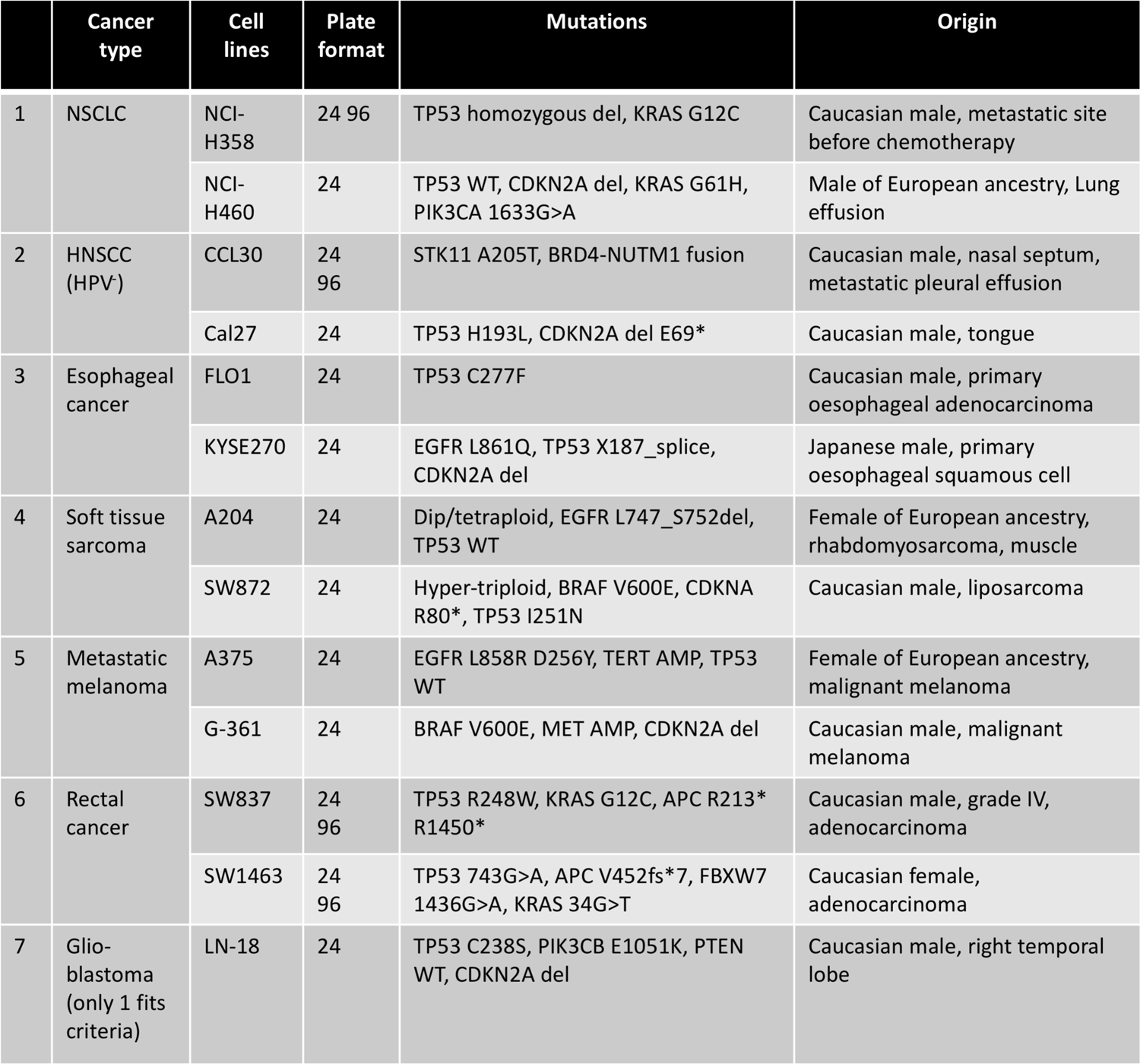
13 cell lines that are amenable to miniaturization. Cancer type and available information on mutation status and patient origin are shown. NSCLC = Non-Small Cell Lung Cancer; HNSCC = Head and Neck Squamous Cell Carcinoma. The information is compiled from (*32-37*) and the following online sources: https://www.cbioportal.org/ https://www.atcc.org/ https://seandavi.github.io/UCCCReporter/reference/cell_line_ancestry.html https://web.expasy.org/cellosaurus/

### Confirmation of equivalent clonogenic chemo-radiation response regardless of plate format

The goal of the clonogenic HTS system is to identify compounds that modulate radiation responses. Therefore, it was necessary to confirm that cell lines exhibiting equivalent radiation responses in miniaturized formats also exhibit equivalent chemo-radiation responses. This was assessed using chemotherapeutics that represent standard-of-care for each of four cancer indications, specifically, Cisplatin for HNSCC; 5-FU, Oxaliplatin and Irinotecan for rectal cancer; Dacarbazine for metastatic melanoma; and Temozolomide for glioma. Using clinically relevant doses for each chemotherapeutic and the same 3-dose radiation design as in Figure 1, dose-modifying factors (DMF’s) were calculated using linear-quadratic plots for clonogenic survival for each well format, cell line and appropriate chemo-radiation combinations (*15*). Importantly, the miniaturized clonogenic formats exhibit equivalent chemo-radiation responses, in addition to radiation responses (Figure 2 and Supplemental Fig. 2; ‘equivalent’ defined as averages that are within one standard deviation from each growth condition).

**Figure 2.**
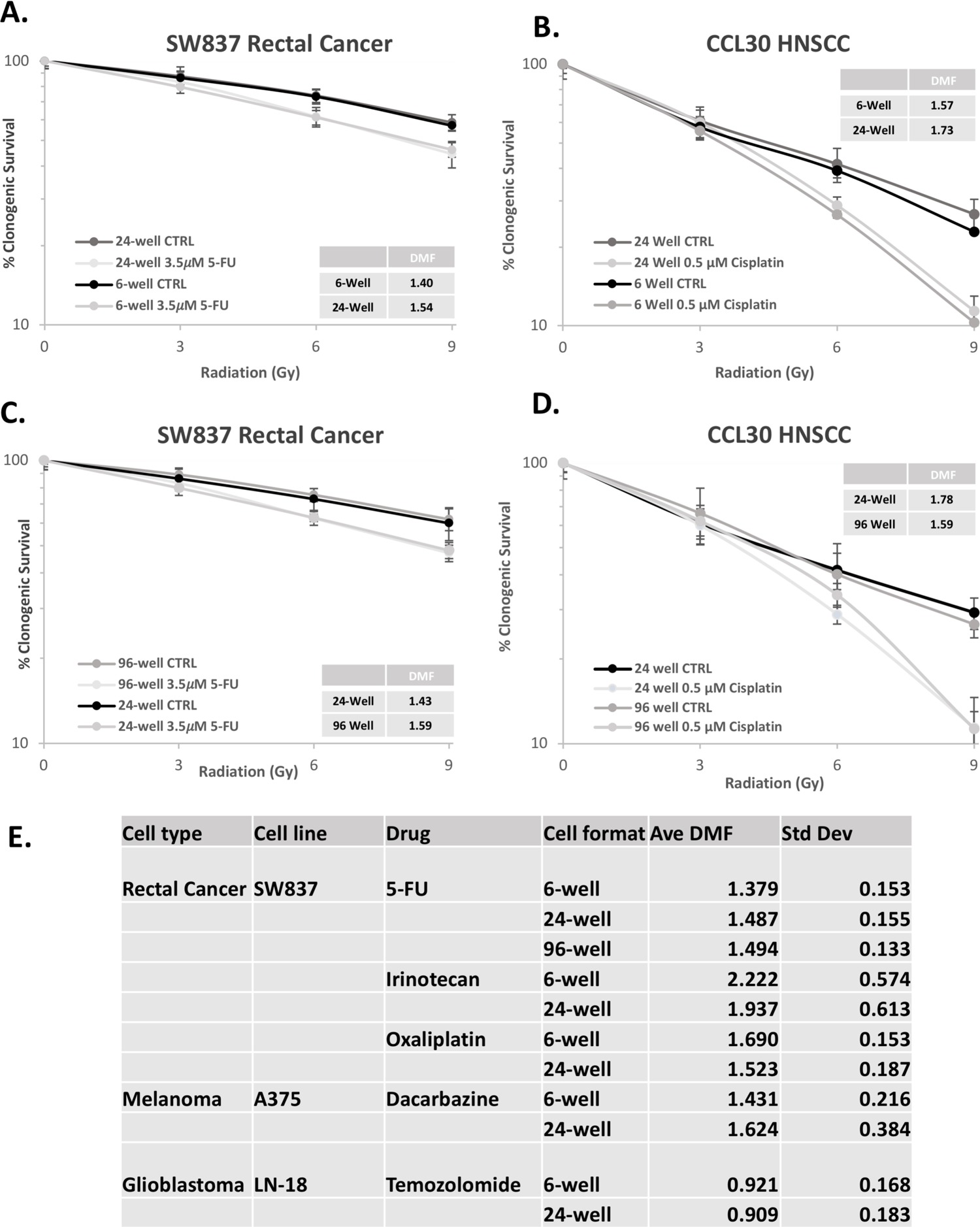
Chemoradiation responses in 6-, 24-and 96-well plates. (A-D) CCL30 HNSCC and SW837 rectal adenocarcinoma cells were seeded at the same relative density (cells/surface area) in 6-well (4 ml/ well), 24-well (1ml/well) and 96-well (150 μl/well) plates and irradiated 24 hours after seeding and drugs were added immediately after irradiation. Clonogenic survival was determined 10 days post radiation. Dose modifying factor (DMF) was computed according to (*15*). (E) Summary of additional known radiation sensitizers with SW837, A375 and LN-18 in different plate formats.

### Automated colony scoring by nuclear position yields equivalent results as manual counting

The most laborious and time-consuming step in clonogenic assays is the counting of colonies, with discrimination between true clones and those that experienced clonogenic death (below the 50-cell/colony threshold). While typically done manually, recent publications describe several automated counting methodologies. A common feature amongst these is the use of area ‘masks’ to identify colonies that have at least 50 cells (Figure 3A, pink circle). While likely accurate under control conditions, this methodology is error-prone in the context of irradiation as radiation can induce cell spreading in some cell lines, thereby changing the area occupied by 50 cells between control and irradiated samples. The extent of cellular spreading is cell type and radiation dose-dependent and can increase the area of a 50-cell colony by as much as 50% (Figure 3B). While this problem may be partially addressed by quantifying the area of a 50-cell colony at different radiation doses and using masks of differing sizes, this methodology would be further confounded by morphological changes induced by drug and/or drug-radiation combinations in a compound library screen. A robust HTS system must accommodate these variables; therefore, we employed an alternative method.

**Figure 3.**
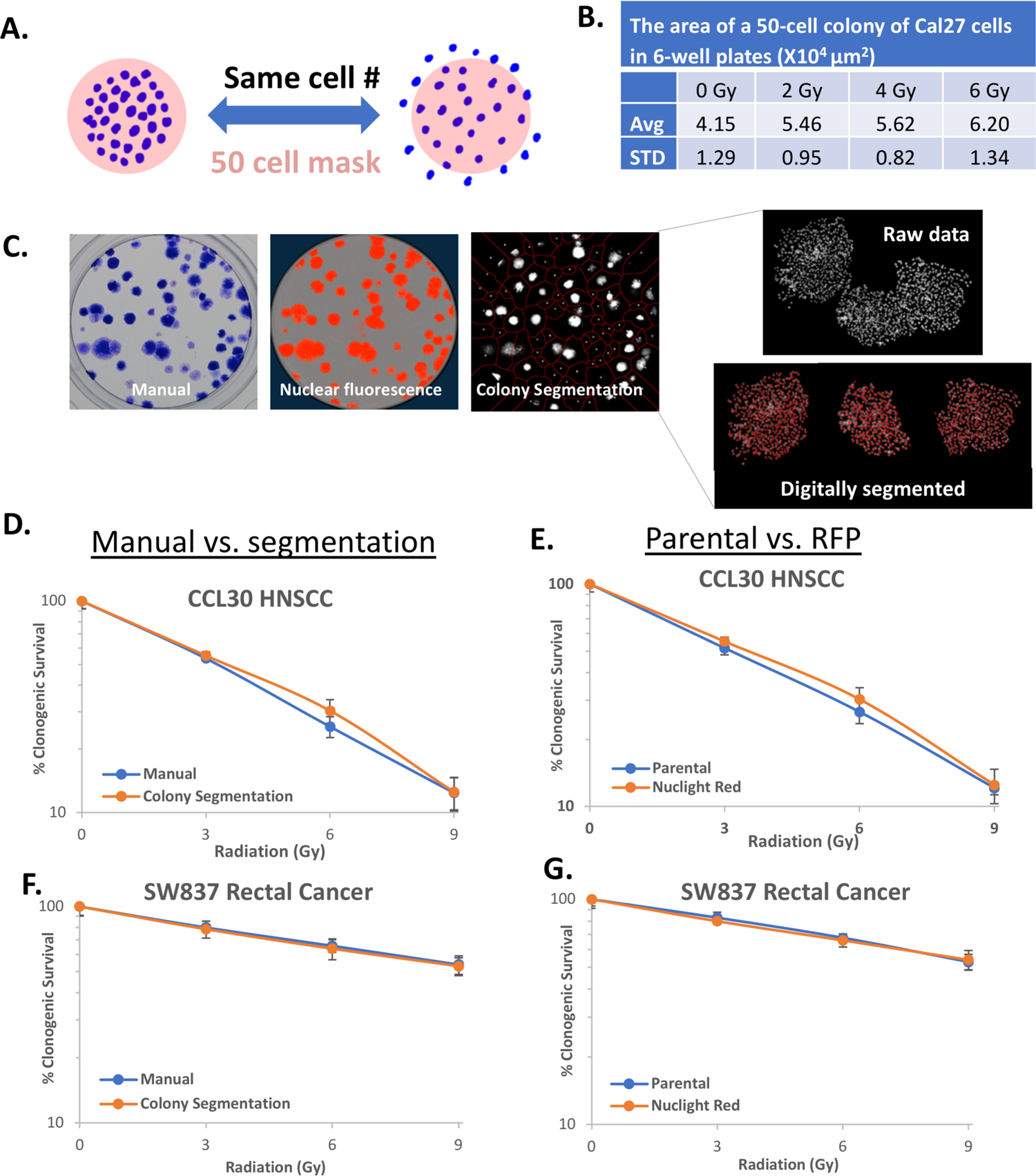
Colony segmentation allows automated colony counting. (A) A diagram to illustrate why a constant area mask cannot be used when cells spread after irradiation. (B) The area of a 50-cell colony increases with radiation for Cal27 HNSCC cells. (C) The same well of a 6-well plate is shown stained for manual counting or imaged for nuclear RFP. ‘Colony segmentation’ panel shows the RFP image after automated digital segmentation. Three colonies are shown magnified to illustrate how the raw nuclear images are grouped or ‘segmented’ into separate colonies based on the position of the nuclei. (D-G) Manual counting and automated segmentation show similar results in clonogenic survival after irradiation as do parental and RFP-expressing derivatives of CCL30 HNSCC and SW837 rectal cancer cells.

We developed a nuclear fluorescence-based, automated colony counting method that utilizes cell line specific parameters to segment colonies and individual nuclei from images of live cells. It is comprised of three steps: image preprocessing to eliminate background and spurious fluorescence, colony segmentation (Figure 3C), and nuclei segmentation to determine nuclear number for each colony (see Methods for details). A delimited text file is generated containing each colony’s relative position within the well along with the number of nuclei in it. A 50-nuclei threshold then identifies clonogens. Fluorescent nuclear dyes are amenable to this methodology but, we found that these have poor reproducibility, low signal to noise ratios, and require stain/de-stain steps (not shown). Instead, we generated stable nuclear RFP (Nuclight Red)-expressing variants for cell lines in Table 1 and validated their use by fixing and staining as in traditional clonogenic assays after live fluorescence images have been acquired (Figure 3C). This allowed us to compare directly automated to manual counts for the same wells, which were found to be equivalent (Figure 3D,F). Survival curves were also equivalent between parental and nuclear RFP variants, suggesting that stable transfection of nuclear RFP did not alter radiation responses (Figure 3E,G). Similar results were observed for all cell lines in Table 1 (Supplemental Figure 3) but we chose to use CCL-30 and SW837 in pilot screens because of their robust growth, well-defined and compact clonogens, and amenability for 96-well miniaturization (Figure 1A-B, D-E and Supplemental Fig. 2).

### Automation of the clonogenic HTS screen

With cell lines amenable to clonogenic miniaturization identified and a validated methodology for automated clonogenic counting in place, we next developed a customized instrument that integrates automated cell- and liquid-handling, irradiation with dosimetry monitoring, and cell plate incubation (Supplemental Figure 4A). Automated seeding of SW837 cells at clonogenic density and radiation survival following RT is consistent across 96-well plates (Supplemental Figure 4B-C). Clonogenic survival with varying drug treatment volumes up to 25 μl is consistent, providing significant flexibility for HTS drug administration (Supplemental Fig 4D). Clonogenic survival is unaffected at up to 0.75% DMSO (drug solvent), more than an order of magnitude greater than the concentration expected to be used in the clonogenic HTS (Supplemental Fig 4E). Similar results were observed when using CCL-30 cells (data not shown). Finally, radiation dosing as measured via robotically placed nanoDot (Landauer) dosimetry is consistent amongst plates (for example, 4.118±0.096 Gy received by 6 plates irradiated with 4 Gy).

As stated previously, CCL-30 and SW837 cells were chosen for pilot clonogenic HTS screens, with the overall goal to identify compounds that modulate clonogenic survival following RT. Therefore, for the primary screen radiation doses that induce partial growth inhibition (30-40%; 3 Gy for CCL-30, 9Gy for SW837) were chosen as this provides conditions for identifying modulators that increase or decrease clonogenic radiation survival. With these conditions, we performed the clonogenic HTS in each cell line with the 1584 compound Diversity Set VI library from the Developmental Therapeutics Program of the National Cancer Institute. The Diversity Set library is assembled from a larger (140,000) molecule library by grouping chemicals according to structure and selecting representative members. Thus, we would be sampling many structurally diverse chemical families with this library.

### Robustness of the clonogenic HTS screen

Cells were seeded in 150 μl, irradiated 24 hr post-seeding, and then treated to a final drug concentration of 1 μM in total volume of 175 μl, with a final DMSO concentration of 0.01%. Cells were seeded to average clonogenic densities of 66.0±7.7 (CCL30) and 110.2±13.2 (SW837) per well. Each screen was conducted in seventeen 96-well plates, with a minimum of six vehicle (DMSO) controls per plate. Live fluorescent images were obtained 10-days post-treatment with an IncuCyte S3. Automated clonogenic segmentation was performed for each cell line in batch, and average growth and standard deviations (Z-score) for each cell line were calculated (Figure 4A-B). To assess the dynamic range of the primary screen, we examined the responses between controls and compounds exhibiting significant growth effects. Vehicle (DMSO) wells function as negative controls, while for positive controls we selected compounds that are 4 (CCL-30) or 4.5 (SW837) Z-scores below the mean. Z’-factor was computed from the average (μ) and standard deviation (σ) of the positive (+) and negative (-) controls, according to the formula Z’ = 1-[(3σ_c+_ + 3 σ_c-_)/Iμ_c+_ - μ_c-_I] exceed 0.5 in both cell lines (Figure 4C), indicative of an HTS with ‘excellent’ robustness and dynamic range (*16*).

**Figure 4.**
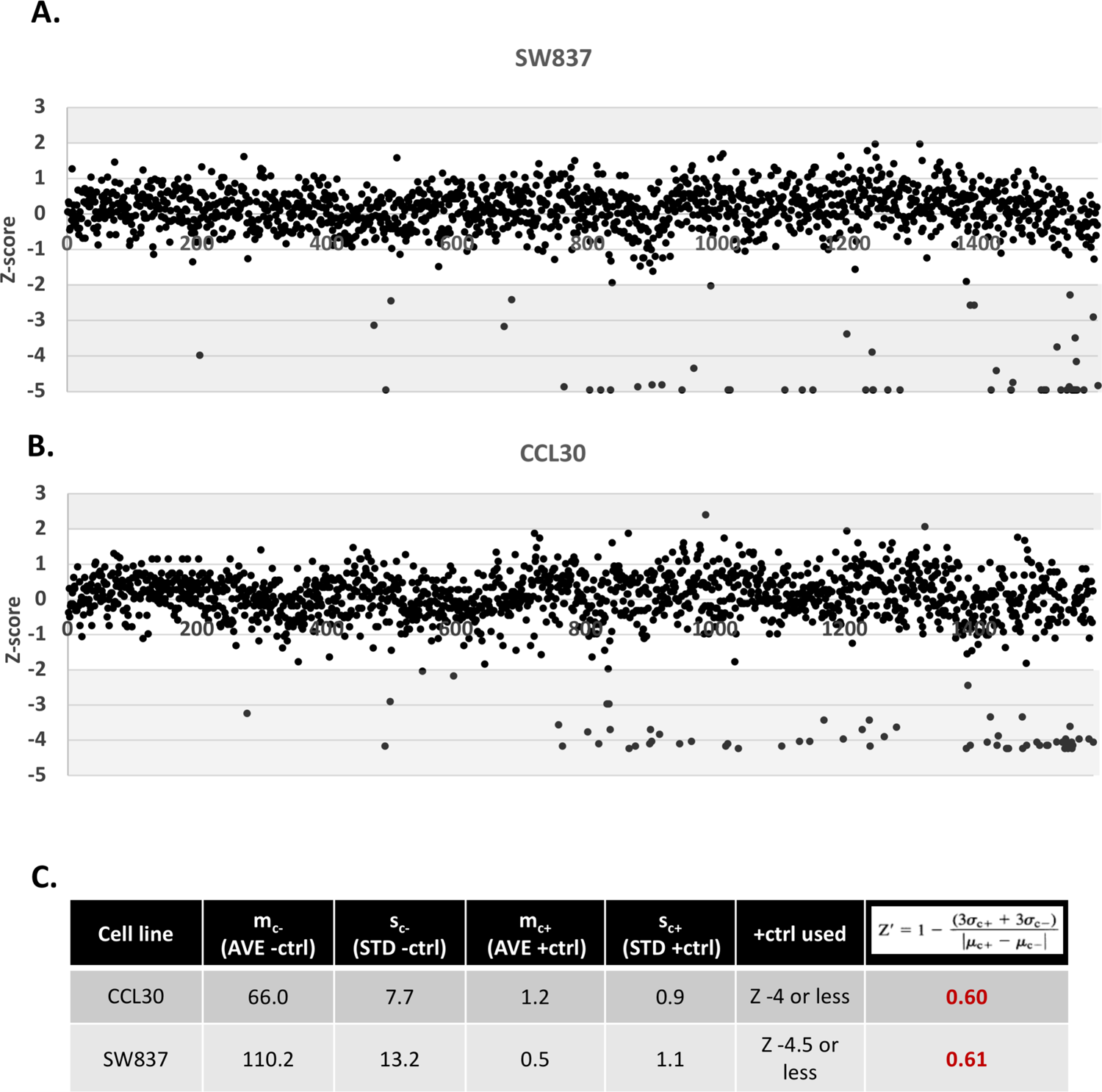
Results of NCI Diversity Set VI screen for two cell lines. (A-B) Data from the pilot screen through the Diversity Set of compounds using SW837 (A) and CCL30 (B) cells. For both cell lines, the data for individual drugs are arranged along the x-axis in same order that they occur on the plates of the drug library. (C) Screen parameters and Z’ table.

### Reproducibility of the clonogenic HTS hits

A common threshold for calling hits in robust HTS screens are effects greater than 2 standard deviations from the mean (*17*). In our primary clonogenic HTS screen, 54 compounds in SW837 and 62 compounds in CCL-30 increased the efficacy of radiation by greater than two standard deviations (>2 Z-score), while no compounds for either cell line exhibited significant radiation protective or growth enhancement effects (Figure 4A-B, shaded areas). Importantly, 41 compound hits were shared between the two cell lines (Figure 5A). To access reproducibility of the identified shared hits, we retested each under the same conditions as the primary screen (1 μM drug plus 3Gy in CCL-30, or 9Gy in SW837). Since the reproducibility test does not include the rest of the library as a control, we assessed growth inhibition for both screens relative to vehicle (DMSO) controls. 39 of the 41 hits (95%) reproducibly exhibited growth inhibition greater than 2 standard deviations from the control in both cell lines (Figure 5B-C). The two compounds that did not reproduce were shared between the cell lines and exhibited no activity in the re-test. Based upon these data we conclude that the clonogenic HTS is highly reproducible.

**Figure 5.**
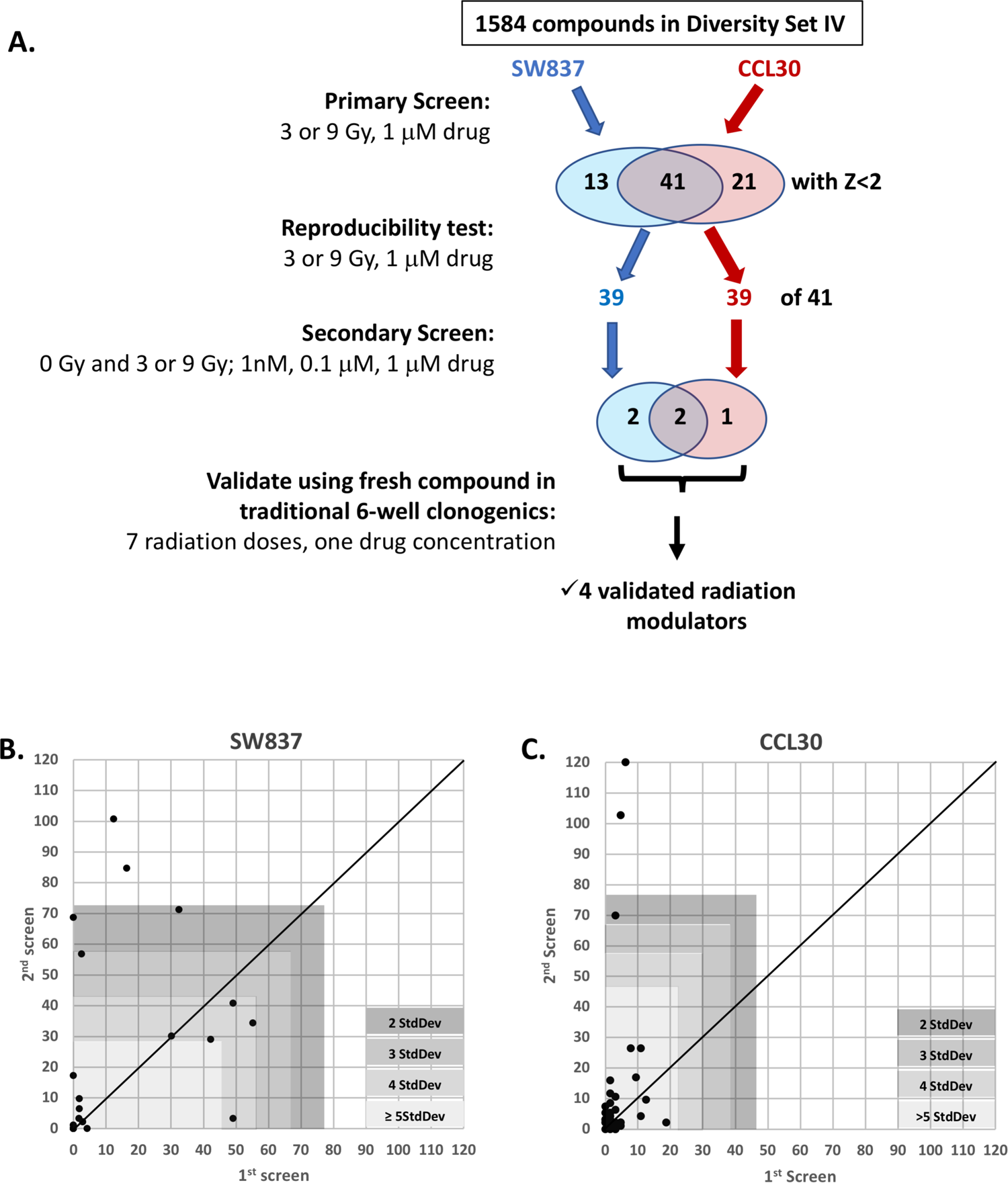
Results of NCI Diversity Set VI screen for two cell lines. (A-B) Retest of 41 hits shared between the two cell lines from the Diversity set screen. Percent (%) clonogenic survival relative to DMSO vehicle controls from the pilot screen (1st screen) are plotted against the same values in the re-test (2nd screen) for SW837 (A) and CCL30 (B). Thirty nine of 41 were within two STD from the vehicle average in both screens, demonstrating reproducibility.Note that the data point at the coordinates 0,0 is comprised of multiple over-laid points; these are wells with no growth in both screens. (C) A summary of screening steps and the results.

### A secondary screen differentiates radiation modulators from single agent activity alone

The clonogenic HTS screen was performed on irradiated cells, and it is therefore expected that some hits will result from robust single agent activity alone rather than radiation modulatory effects. Furthermore, radiation modulation could result in a simple sum of the drug and radiation effects (additive interaction) or greater than the simple sum (synergistic interaction). To identify true radiation modulators in the 39-compound hit pool, each compound was retested with and without radiation (3Gy for CCL-30, 9Gy for SW837) at varying concentrations (10 nM, 0.1 μM, and 1 μM). In SW837 cells, 6 compounds exhibited potent single agent activity, producing growth inhibition of greater than 90% at 10 nM drug (for example NSC330770 in Supplemental Figure 5A). 29 compounds exhibited what is expected if combination with radiation is additive (for example, NSC250594 in Supplemental Figure 5B). And 4 compounds exhibited greater than additive interaction with radiation, which is what is expected of true radiation modulators (Figure 6A). Two of these compounds (NSC177365 and NSC228155) exhibited greater than additive activity at 1 μM, the drug concentration used in the primary screen. The other two compounds exhibited complete single agent growth inhibition at 1 μM, but partial single agent activity when reduced to 10 nM (NSC94600 and NSC317003). And at 10 nM both showed greater than additive effects when combined with radiation. Similar results were obtained with CCL30 cells; three of 39 compounds showing greater than additive effects with radiation. All three showed robust single agent activity at 1 μM, with radiation modulation becoming apparent at 1-100 nM depending on the compound (Figure 6B and Supplemental Table 1A). Of these hits, two are shared between both cell lines, resulting in a total of 5 potential radiation modulators identified (Supplemental Table 1B).

**Figure 6.**
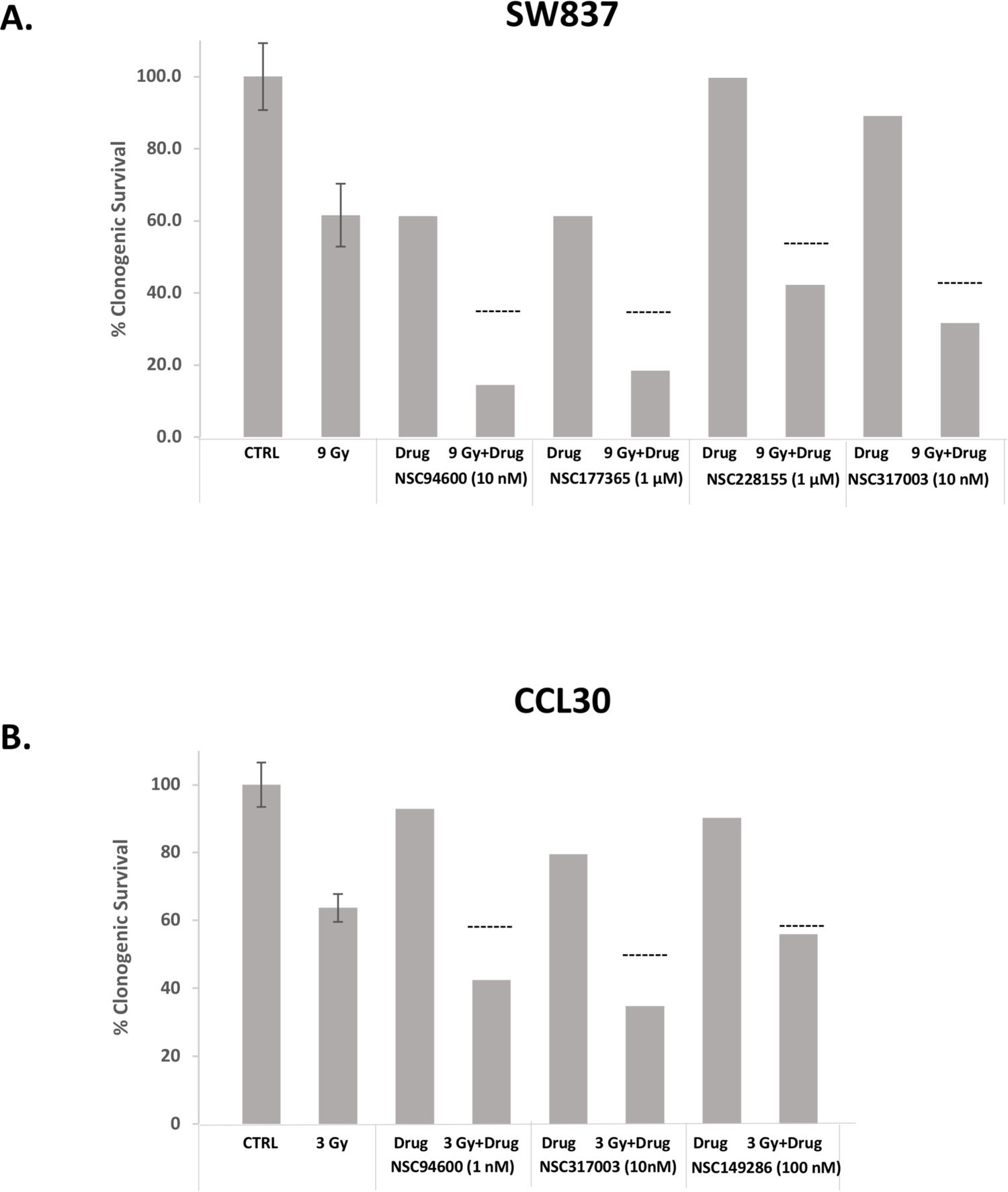
A secondary screen differentiates radiation modulators from cytotoxic agents. (A-B) Clonogenic survival after treatment with DMSO (control), radiation alone, drug alone or the combination with drug with radiation in SW837 (A) and CCL30 (B) cells. Dashed lines show the expected survival if the drug and radiation acted in an additive manner. Survival below the dashed lines of combined treatments are likely caused by a synergistic interaction between the drug and radiation. Control and radiation alone samples are an average of 6 replicate wells each. The others are from one well per condition.

### Validation of pilot HTS hits in traditional clonogenic assays

To this point, the data from the primary clonogenic HTS and confirmatory experiments were obtained using compounds supplied as 10 mM DMSO stocks in multi-well plates from the NCI. Our previous experience indicates that plated compounds in solution can exhibit reduced activity relative to freshly acquired and prepared solutions of the same compound. Therefore, we acquired from the NCI vialed solid aliquots of the five potential radiation modulators and prepared fresh DMSO stocks. As expected, freshly prepared compounds show varying degrees of activity as single agents relative to plated drug, ranging from similar (NSC94600, NSC317003, NSC149286) to 10-fold more active (NSC177365, NSC228155) (Supplemental Figure 5C). Therefore, to test for radiation modulation, we titrated each fresh compound to induce 25-30% growth inhibition. We then used fresh dilutions at these respective concentrations to generate survival curves with a broad range of 9 radiation doses, and calculated the resulting DMF for each hit. These experiments were performed first in 96-well plates as with the clonogenic HTS (Supplemental Figure 6), and then validated with traditional 6-well clonogenic assays (Figure 7).

**Figure 7.**
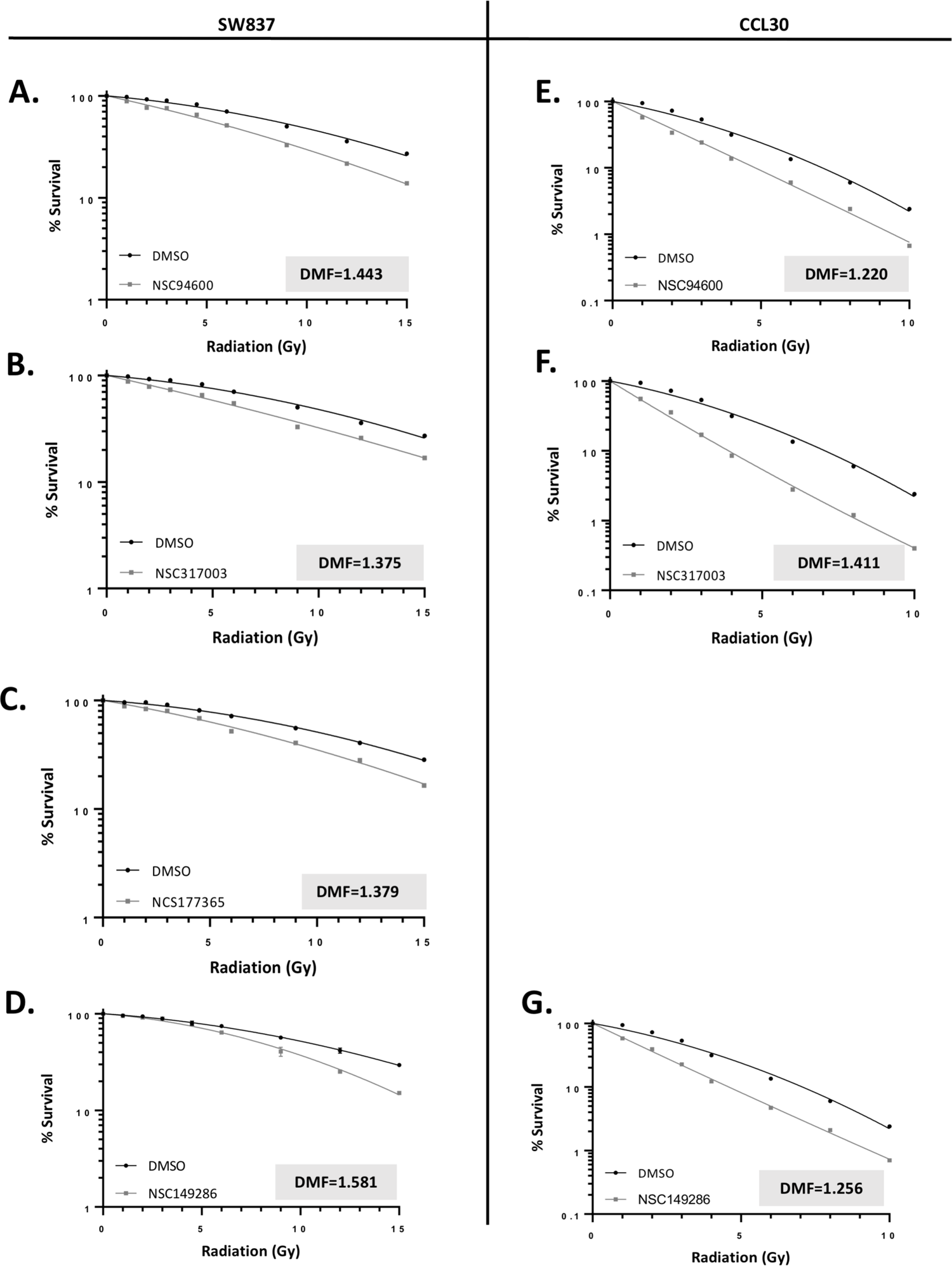
Radiation modulators found in the HTS were validated in 6-well clonogenic assays. (A-D) Four compounds that showed greater than additive effect with radiation in the secondary screen using SW837 cells were assessed in traditional 6-well clonogenic assays using 8 radiation doses. Dose Modifying Factors indicate that four enhanced the effect of radiation. Drug concentrations used with SW837 cells were 8 nM (NSC94600), 15 nM (NSC317003), 250 nM (NSC 177365) and 150 nM (NSC149286). Drug concentrations used with CCL-30 cells were 1 nM (NSC94600), 10 nM (NSC317003) and 100 nM (NSC149286). (E-G) Three compounds that showed greater than additive effect with radiation in the secondary screen using CCL30 cells were assessed in traditional 6-well clonogenic assays using 7 radiation doses. Dose Modifying Factors indicate that three enhanced the effect of radiation.

In the 96-well format, three of four hits found in the SW837 screen recapitulated radiation modification with DMF’s >1.4 (Supplemental Figure 6A-C), with the exception being NSC228155 (not shown, see Discussion). In CCL-30 cells, all three hits recapitulated (Supplemental Figure 6E-G). Interestingly, NSC149286, a hit in CCL-30 screen but not SW837 screen, was found also to modulate radiation in SW837 when tested with 9 radiation doses; a possible explanation for this result is provided in the Discussion. More important, all compounds that exhibited radiation dose modification in miniaturized formats exhibited nearly identical modification in traditional formats, with DMFs ranging from 1.375 to 1.581 in SW837 (Figure 7A-D) and from 1.220 to 1.411 in CCL-30 (Figure 7E-G). We conclude that the clonogenic HTS methodology can robustly identify compounds that function as radiation modulators in traditional clonogenic formats, and therefore can serve as a functional novel drug discovery platform with a true clonogenic endpoint.

## Discussion

### A validated HTS screen with a clonogenic endpoint

We describe here an integrated system capable of performing HTS screens with a clonogenic endpoint to identify radiation modulators. This system differs significantly from previously described methodologies in key aspects: 1) while miniaturized, this is a true clonogenic assay wherein cells are seeded at clonogenic densities and grown over traditional time periods, with traditional endpoints and statistical practice; 2) the ability to perform clonogenic HTS in multiple cell lines for seven cancer indications where RT is a standard treatment module; 3) automated colony scoring based upon nuclei count rather than colony size masks, ensuring accurate colony counts regardless of radiation/drug induced changes in colony morphology; and 4) integrated radiation dosimetry ensuring consistency over HTS runs. Our data demonstrate that this clonogenic HTS system can identify compounds that exhibit robust radiation Dose Modifying Factors (DMF) in traditional clonogenic assays. Throughput is markedly increased due to miniaturization and automation of colony counting. The significance of these innovations cannot be overstated: the traditional clonogenic assay is laborious, including manual cell seeding, irradiation, drug treatment, fixing and staining, that ultimately culminates in exceedingly tedious manual counting of colonies. Simple human limitations reduce screening capability to no more than a dozen compounds per day. In contrast, our automated clonogenic HTS is capable of processing over 2000 compounds a day, with throughput capacity that can be scaled linearly with additional incubator space, off-the-shelf imaging units, and computer processing.

### Clonogenic assay harmonization

There is increasing recognition that disparate clonogenic assay protocols make comparing data across research labs nearly impossible (*5, 18*). Worse, some agents identified as radiation sensitizers appear not to be so, and vice versa depending upon the research group and assay protocol used (*14*). The clonogenic HTS described here address several parameters of concern. We assessed the role of plate format (well size) in relation to clonogenic survival and identified conditions wherein clonogenic radiation survival is equivalent between miniaturized and traditional 6-well formats (Figure 1). Our automated colony segmentation methodology controls for radiation induced variations in cell and colony shape/size by assessing nuclear position/number rather than size masks (Figure 3). The resulting positional identification of actual nuclear counts allows for the accurate quantification of colonies greater than 50 cells, in agreement with the parameters of the original clonogenic assay colony (*6*). We standardized protocols to achieve a minimum colony number of 50 per well, as indicated by the literature (*13*), and exceed this parameter in the pilot screens (Figure 4C, the second column). We validate candidate radiation modulators with a broad range of radiation doses (9 total), allowing for monitoring of the initial “shoulder” and exponential portions of survival curves (Figure 7 and Supplemental Figure 6). We use DMF to quantify radiation modulation here but have developed automated programs capable of computing additional parameters including, survival fraction at a given radiation dose and α/β ratio from linear-quadratic plots. Finally, we address inter-person variation by fully automating the clonogenic process, and conducting the pilot screens independently with two researchers, in two different cell lines.

### Radiation modulators identified in the pilot screen

The pilot screen and reproducibility tests identified five potential radiation modulators (Figure 5A): two shared between the SW837 and CCL-30 screens (NSC94600, NSC317003), two exclusive to SW837 (NSC177365, NSC228115), and one exclusive to CCL-30 (NSC149286). Of these, only NSC228155, an EGFR agonist (*19*), did not fully validate through subsequent experiments. Specifically, radiation modulation was reproducibly observed with plated drug (Figure 6A), however fresh compound exhibited 10-fold greater single agent activity (Supplemental Figure 5C) and showed no modulation under any condition tested. The other four exhibited reproducible radiation modulation in traditional 6-well clonogenic formats, regardless of variables between plated and fresh drug single agent activity. Available data on these four compounds illustrate how our clonogenic HTS can identify known as well as new radiation modulators.

One hit shared between the cell lines is Camptothecin (CPT, NSC94600) a well described DNA intercalator and potent topoisomerase I inhibitor and a known radiation sensitizer (*20*). Clinically relevant derivatives of Camptothecin include Irinotecan (Camptosar), a known radiation modulator and standard of care for colorectal cancers which SW837 cells represent (*20, 21*). Our clonogenic HTS was performed blind, using a library based on chemical structure and not for anti-cancer activity. Therefore, identification of CPT is proof-of-concept that the screen is capable of accurately identifying compounds that modulate radiation effects.

NSC317003, was shared between cell lines and is a structural analog of lucanthone and its metabolite hycanthone (Supplemental Figure 8A). Lucanthone, a known DNA intercalator and topoisomerase I/II inhibitor, was historically used to treat parasitic diseases, but more recently has been studied as a radiation sensitizer (*22*). Lucanthone inhibits post-radiation DNA repair in human cancer cells (*23*), accelerates the regression of brain metastases in patients receiving radiation therapy (*24*), and has been in clinical trials as a radiation sensitizer for the treatment of Glioblastoma (NCT01587144) and brain metastases of NSCLC (NCT02014545). Lucanthone and hycanthone are not in Diversity Set VI but identification of a structural analog in our clonogenic HTS further validates its ability to identify radiation modulators.

NSC149286 is an analog of curcumin (Supplemental Fig. 8B), which is has multiple reported activities including radiosensitization of cancer cells and radio-protection of normal cells [(*25-27*); reviewed in (*28*)]. NSC149286 exhibited slightly greater than additive effect with radiation in CCL-30 cells (Figure 6B) and an additive effect in SW837 cells (not shown). Fresh compound, however, showed robust radiation modulation in both cell lines when tested with 7 radiation doses (Figure 7D and G). The survival curves of SW837 suggest an explanation for the discrepancy between the secondary screen and radiation dose range response data. Specifically, survival with or without drug was equivalent at low doses of radiation, began to diverge at 9Gy, and increased significantly at higher radiation doses (Figure 7D, Supplemental Fig. 6D). The level of divergence at 9Gy in the secondary screen was considered to be within error and therefore scored as additive for SW837. Greater divergence at >9Gy, however, resulted in a DMF>1. These data suggest that while use of a single radiation dose in the pilot and secondary screens maximizes the number of compounds assessed, we may miss modulators that show effect at other doses (false negatives). We believe this tradeoff to be acceptable as compounds that modulate RT under many conditions are a preferred outcome. While curcumin is not in the NCI Diversity Set VI library, identification of a closely related analog further validates the ability of the clonogenic HTS to identify radiation modulators.

The pilot screens also identified a compound with known antineoplastic activity but previously undescribed radiation modulation effects. This hit, specific to SW837 cells, was NSC177365 which has been demonstrated to partially restore folding of the G-quadruplex in human telomerase reverse transcriptase (hTERT) promoters, thereby silencing hTERT transcription and leading to cell death in melanoma cells (*29*). Further, NSC177365 and related compounds have been demonstrated to inhibit cystathionine β-synthase, reducing intratumor production of hydrogen sulfide which plays important roles in cancer proliferation and migration [reviewed in (*30*)]. Finally, a patent for use of NSC177365 as a bone morphogenetic protein (BMP) signaling modulator has been granted (US-10434220-B2), which has direct relation to cancer therapeutics [reviewed in (*31*)]. The ability of NSC177365 to modulate radiation in SW837 cells significantly expands its potential role as a cancer therapeutic. Interestingly, NSC177365 exhibited exquisitely robust single agent activity in CCL-30 cells even at the lowest doses tested, to the point that no relevant compound/RT data could be obtained.

### Standard Operating Procedure and looking forward

The streamlined clonogenic HTS procedure includes 4 critical steps: (1) a primary screen with a single LD_30-40_ radiation dose for the cell line and 1 μM of drug library; (2) the use of Z>2 from the population mean to call hits; (3) a secondary screen with 0 and LD_30-40_ Gy and each hit at 10 nM, 0.1 μM and 1 μM; and (4) validation in 6-well plates using fresh compound titrated to achieve 20-30% growth inhibition on its own. We believe this methodology is a major step forward for the unbiased identification of novel radiation modulators, but represents only the initial uses of this technology. Clonogenic HTS may be adapted to screen for compounds that inhibit regrowth of tumors following any standard therapies, as well as identify efficacious pair-wise combinations among existing therapeutic agents. For example, an IC_50_ dose of a single agent may be applied to all wells, much like single dose radiation, and used as the basis to screen for compounds that modulate that activity towards a clonogenic endpoint. Further adaptions to this methodology in development include 3D tumor growth, co-culture with somatic cells to recreate tumor microenvironments, and assays for immune-cell infiltration in the context of irradiation.

## Materials and Methods

### Cell culture

All cell lines were obtained from ATCC (American Type Culture Collection) except for FLO-1, A204 and KYSE270 which were obtained from Sigma-Aldrich. Cell line identities were confirmed by STR fingerprinting at the University of Colorado Cancer Center Cell Technologies Shared Resource. All cell lines were maintained in DMEM with 10% FBS supplemented with Penicillin (50 units/mL) and Streptomycin (50 μg/mL).

### Transduction with nucRFP lentivirus

Cells with low passage number (20 or fewer) were labelled with Nuclight Red (Sartorius) according to manufacturer’s instructions. Briefly, cells were plated to 15-35% confluence and allowed to adhere overnight. Nuclight Red lentivirus was added at 1 multiplicity of infection (MOI)/cell along with Polybrene (8 μg/mL). After 24 hours, media was replaced with fresh media containing 1 μg/mL puromycin. Media with puromycin was relaced every 48 hours until stable clones were selected.

### Plating efficiency, cell seeding and colony formation

Cells were stained with trypan blue (1:5 in PBS) and manually counted with a hematocytometer. Cell populations with less than 10% trypan blue-stained (dead) cells were serially diluted in media and plated in 24 or 96 well plates to determine plating efficiency (PE) and optimum cell seeding for clonogenic assays. PE varied widely among the 13 cell lines (>90% for A375 to <16% for KYSE270) and were consistent in additional assays for each cell line. Colony formation was monitored daily over 7-10 days to determine an endpoint where colonies were still distinguishable.

### Radiation survival curves of parental and nucRFP-labelled cells

Parental and nucRFP cells were plated in 6, 24 or 96 well formats to give equivalent colony densities and irradiated 24 hours later. After 10 days, parental cells were fixed and stained with 0.1% crystal violet or 0.4% Sulforhodamine B for manual counting whereas nucRFP derivatives were imaged live using the IncuCyte S3 for colony segmentation.

### High throughput screen

The SuviCa high throughput system consists of 4 functional instruments: a 37°C CO_2_ incubator (LiCONiC), a Fluent 780 drug distribution workstation (Tecan), a TEMPEST Liquid Handler (FORMULATRIX), and a custom irradiator (Hopewell Designs) calibrated to deliver 3.2 Gy/min. Cells were counted using trypan blue exclusion described above and plated according to previously determined cell seeding (250 cells/well for CCL30, 200 cells for SW837) in 150 μL/well in 96-well plates. To prevent edge effects of prolonged incubation, outside wells were filled with 200 μL sterile H_2_O. Cells were incubated overnight before irradiation, and drugs were added immediately after irradiation.

The NCI Diversity Set VI is arrayed in 96 well plates at 10 mM in 20 μL DMSO. Upon arrival, compounds were partitioned into duplicate plates at 5 μL per drug and stored at −20°C. For all assays, a replicate plate was thawed only once and diluted 1:40 in complete media 1-2 hours before irradiation. 1.4 μL of diluted compound was added to 200 μL of complete media. The Fluent was programmed to add 25 μL/well to give a final concentration of 1 μM in a total volume of 175 μL/well. Drugs were added after irradiation to avoid X-ray damage to the drugs. Plates were incubated for 10 days before imaging in IncuCyte S3.

### Automated nuclear fluorescent based live-cell colony counting

This is comprised of three steps: image preprocessing, colony segmentation, and nuclei segmentation. Parameters for each step are empirically optimized for cell line and growth format specific conditions, then subject to images from a given experiment in batch. Image preprocessing involves reducing spurious signal from culture well walls (Crop Perc.) and defining the maximum and minimum signal values within the image to be analyzed (Thresh min/max). Colony segmentation involves implementing Gaussian blur functions to merge proximal nuclear signals into discreate colonies (Blur Factor), followed by iterated erode functions to further refine Gaussian colonies and remove other spurious signals (erode iter.), finally implementation of colony prominence thresholds and seeded watershedding functions define actual colonies from the pool of putative Gaussian colonies (colony prominence). Cell segmentation involves similar methodologies, wherein for each identified colony individual nuclear signals are defined by prominence thresholds and seeded watershedding functions (cell prominence). Further parameters allow for previewing of defined portions of a given well image, allowing for optimization of the parameters for image processing and colony/cell segmentation. Colony segmentation output for each well is provided in delimited text format (CSV) containing each clonogens relative position within the well, and the nuclei count for all colonies. Nuclei threshold (>50) is then applied to identify clonogens. A JAVA applet employing this methodology can be obtained here: download link (to be available upon manuscript acceptance).

## Acknowledgements

This work was supported by Small Business Innovation Research phase I contract No. HHSN261201700045C and Phase II contract No. 75N910196C00038 from the National Cancer Institute to SuviCa (PIs: TTS and BF). Work in the Su lab is supported by National Institutes of Health grant R35 GM130374 to TTS.

