## Supplemental Data for "A high throughput screen with a clonogenic endpoint to identify radiation modulators of cancer"

### Supplemental Figure 1.

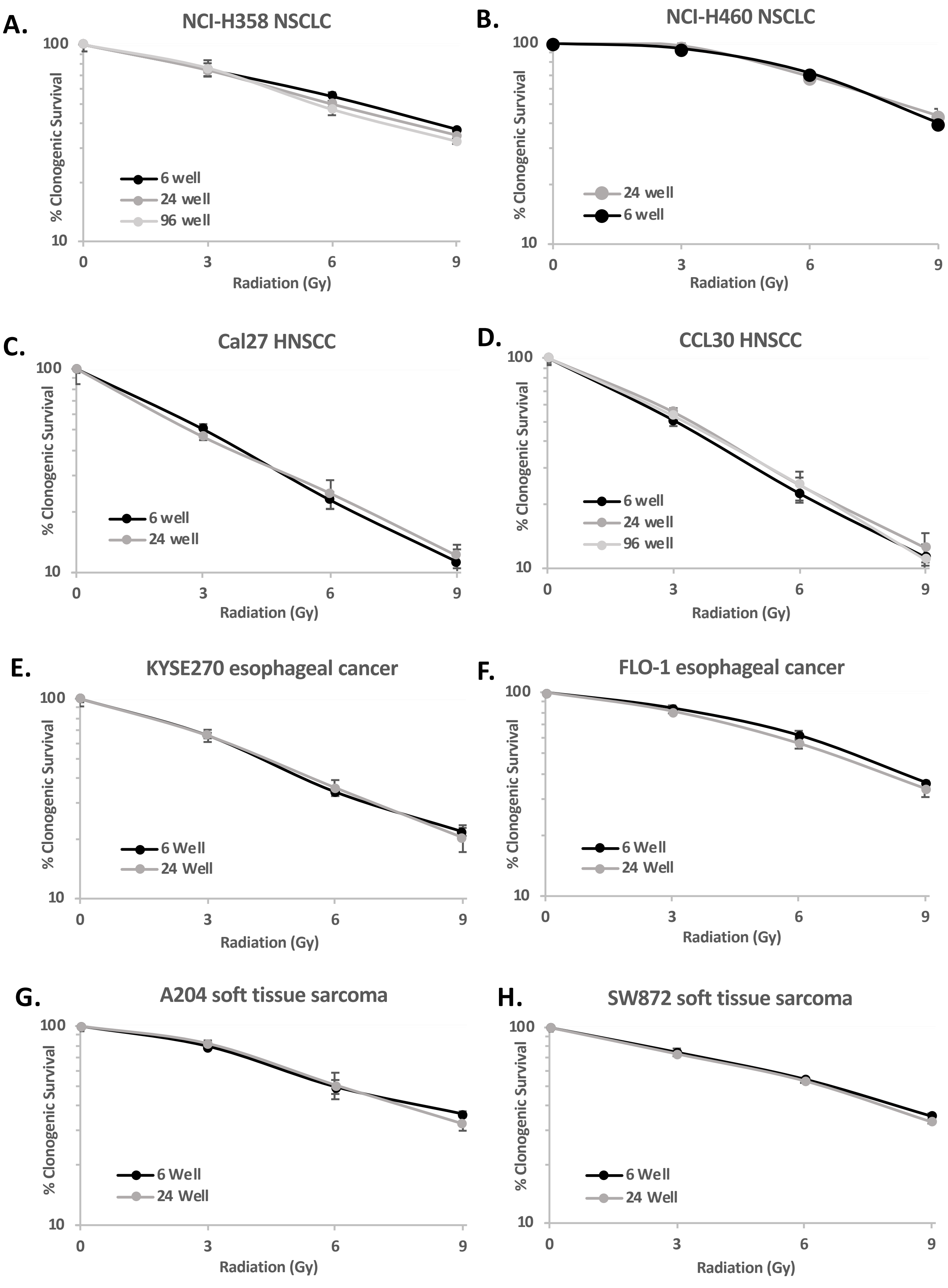

### Supplemental Figure 1 (Continued)

I.

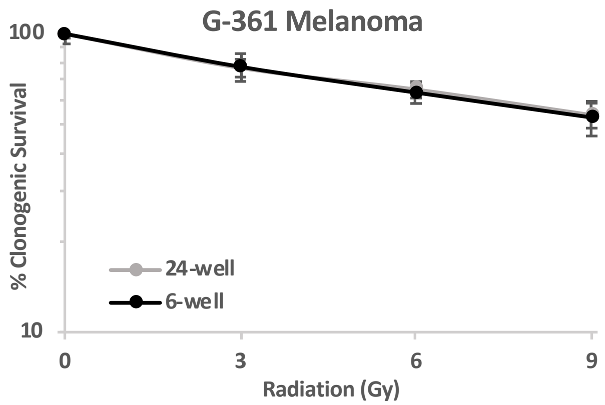


J.

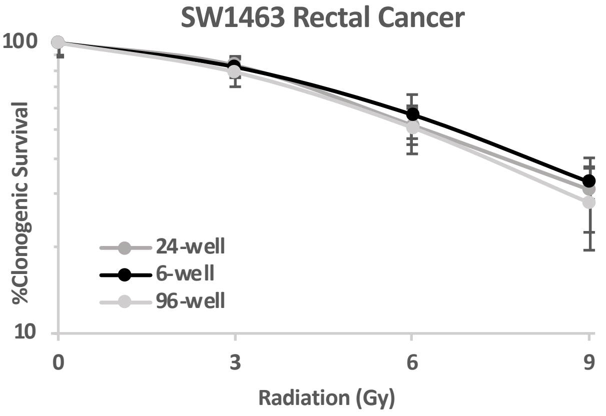


K.

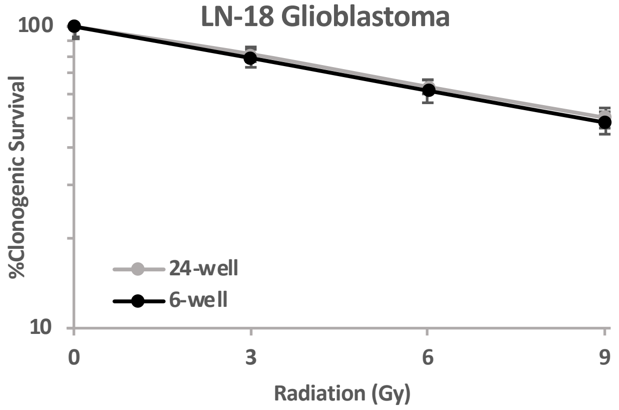


**Supplemental Figure 1**. Additional cell lines that exhibit equivalent radiation response between small well formats and 6-well plates. Cells were seeded in 4 ml (6-well), 1 ml (24-well) or 125 ml (96-well) and irradiated the following day. Growth at a given radiation dose was considered equivalent if average survival was within one standard deviation from the average survival in the 6-well format.

### Supplemental Figure 2.

A

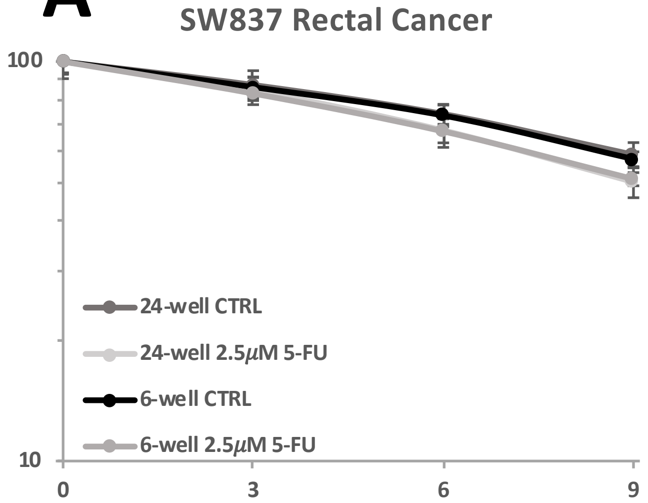


B

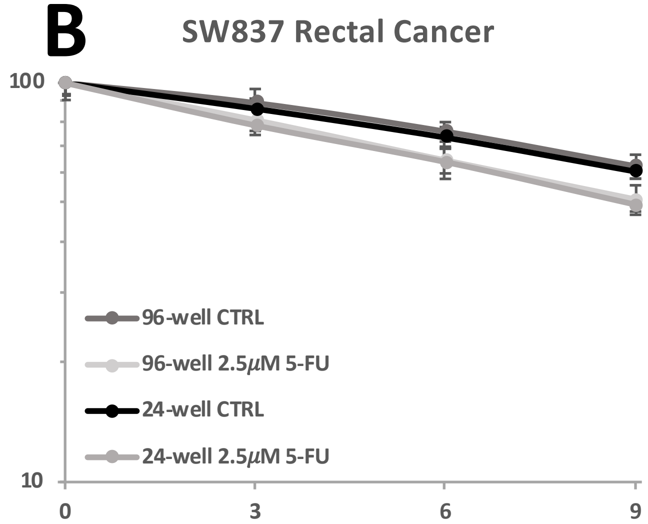


C

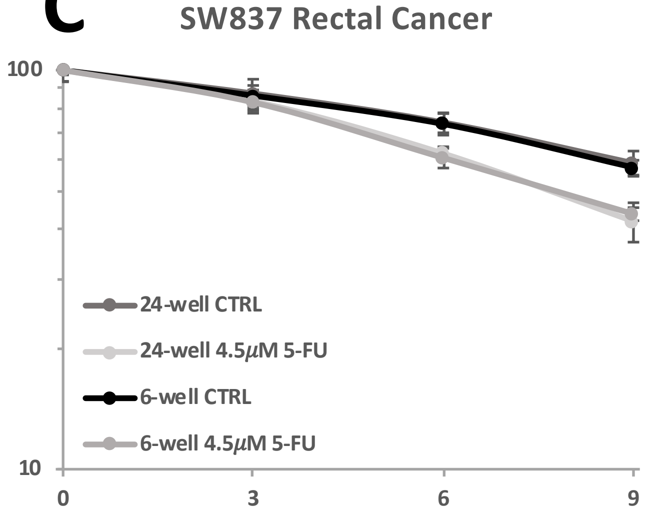


D

| Cell line | Drug | Conc. | Plate format | Average DMF |
|---|---|---|---|---|
| SW837 rectal cancer | 5-FU | 2.5 μM | 6-well | 1.217 |
| | | | 24-well | 1.332 |
| | | | 96-well | 1.400 |
| | | 3.5 μM | 6-well | 1.400 |
| | | | 24-well | 1.487 |
| | | | 96-well | 1.588 |
| | | 4.5 μM | 6-well | 1.520 |
| | | | 24-well | 1.642 |
| | Oxaliplatin | 0.25 μM | 6-well | 1.582 |
| | | | 24-well | 1.391 |
| | | 0.40 μM | 6-well | 1.799 |
| | | | 24-well | 1.655 |
| | Oxaliplatin | 0.25 μM | 6-well | 1.816 |
| | | | 24-well | 1.504 |
| | | 0.40 μM | 6-well | 2.628 |
| | | | 24-well | 2.370 |
| A375 melanoma | Dacarbazine | 100 μM | 6-well | 1.199 |
| | | | 24-well | 1.278 |
| | | 150 μM | 6-well | 1.469 |
| | | | 24-well | 1.557 |
| | | 200 μM | 6-well | 1.625 |
| | | | 24-well | 2.037 |
| LN-18 glioblastoma | Temozolomide (TMZ) | 100 μM | 6-well | 1.012 |
| | | | 24-well | 0.977 |
| | | 200 μM | 6-well | 1.025 |
| | | | 24-well | 1.048 |
| | | 400 μM | 6-well | 0.727 |
| | | | 24-well | 0.702 |

**Supplemental Figure 2**. Chemo-radiation responses in 6-, 24- and 96-well plates for additional cell lines.
(A-C) Primary data for SW837 cells at different 5-FU concentrations and different plate formats are shown.
(D) DMF was computed as described for Figure 2. DMF values are functionally equivalent for each drug concentration across different plate formats.

#### Supplemental Figure 3.

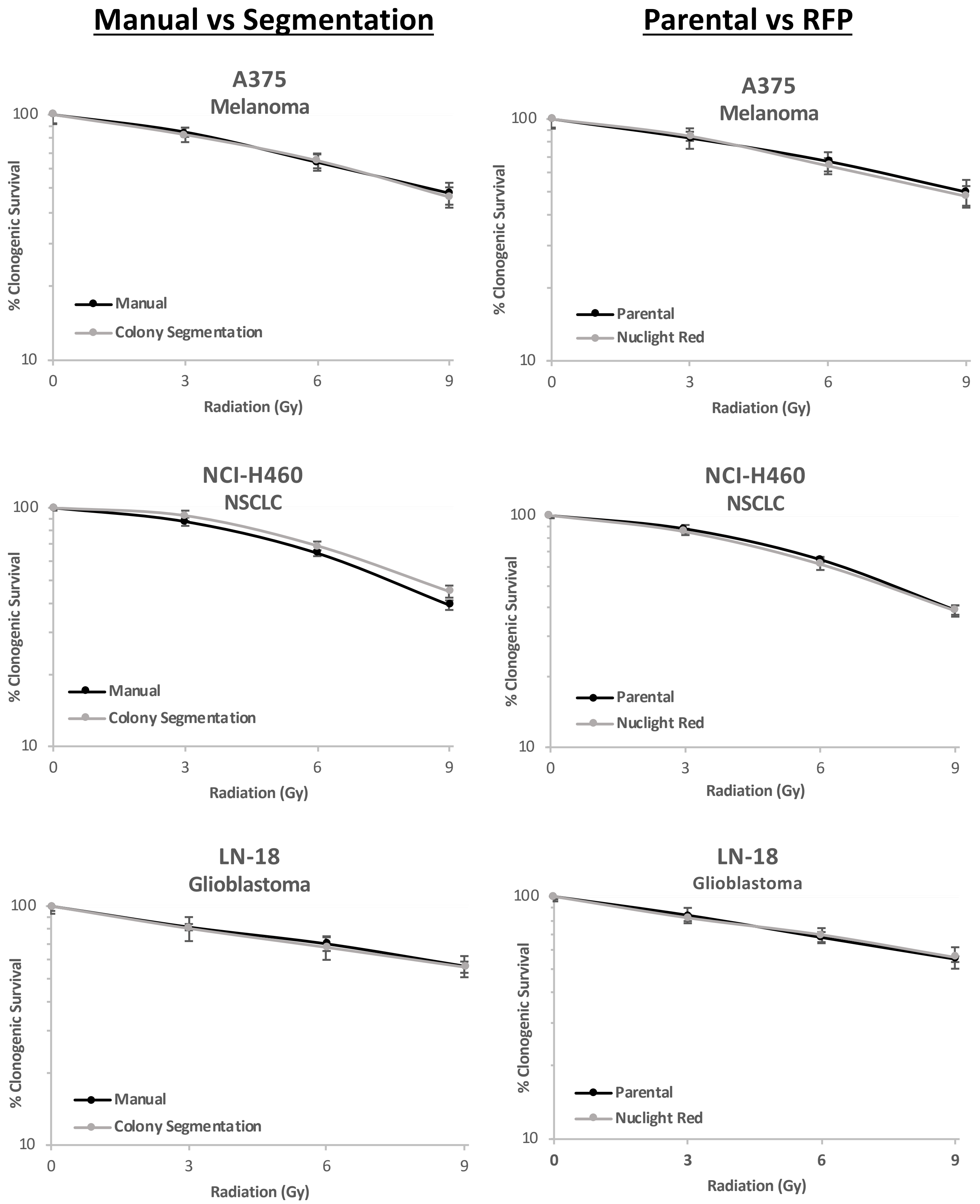


**Supplemental Figure 3**. Comparison of manual counting vs. automated segmentation and parental vs. nuclear RFP derivatives, for three additional cell lines.

#### Supplemental Figure 4.

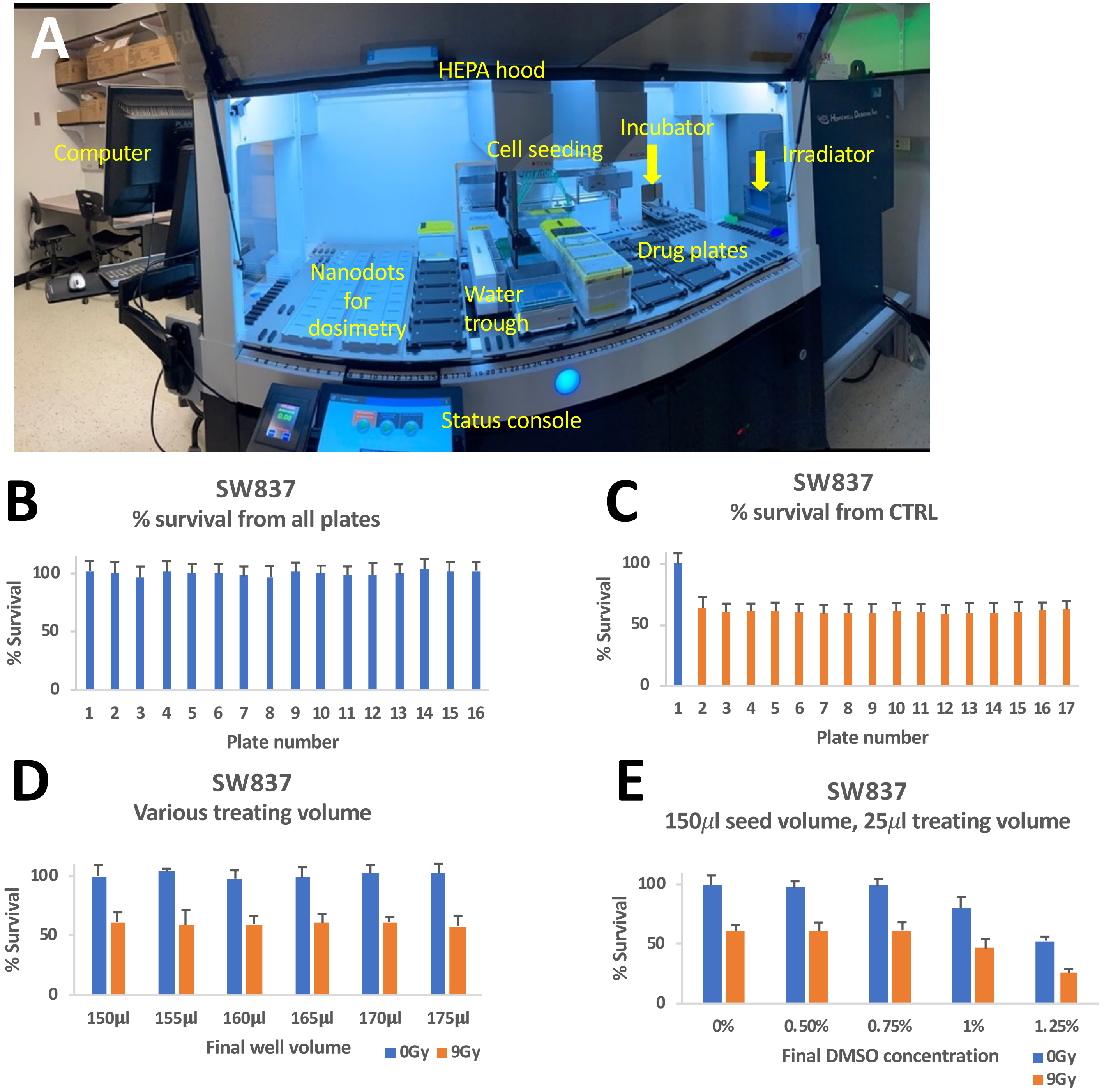


**Supplemental Figure 4. A high throughput workstation for clonogenic assays.**

(A) The clonogenic HTS instrument. Cell/liquid handling, dosimetry, radiation and incubation are integrated into a single system.

(B-C) Quality control for automated plating and irradiation. Clonogenic survival without irradiation (B) and after irradiation with 9 Gy (C) was consistent across multiple plates of cell seeding. Survival was normalized to the average from plate 1.

(D) Clonogenic survival with or without irradiation was consistent across different final volumes after mock drug-addition in 96-well plates. 175 ml final volume (150 to seed + 25 to add drug) was used in the pilot screen.

(E) Final DMSO concentration of up to 0.75% v/v did not affect clonogenic survival with or without irradiation. Final DMSO concentration of 0.01% was used in all subsequent experiments. The data are average of 6 technical replicates in B-E.

#### Supplemental Figure 5.

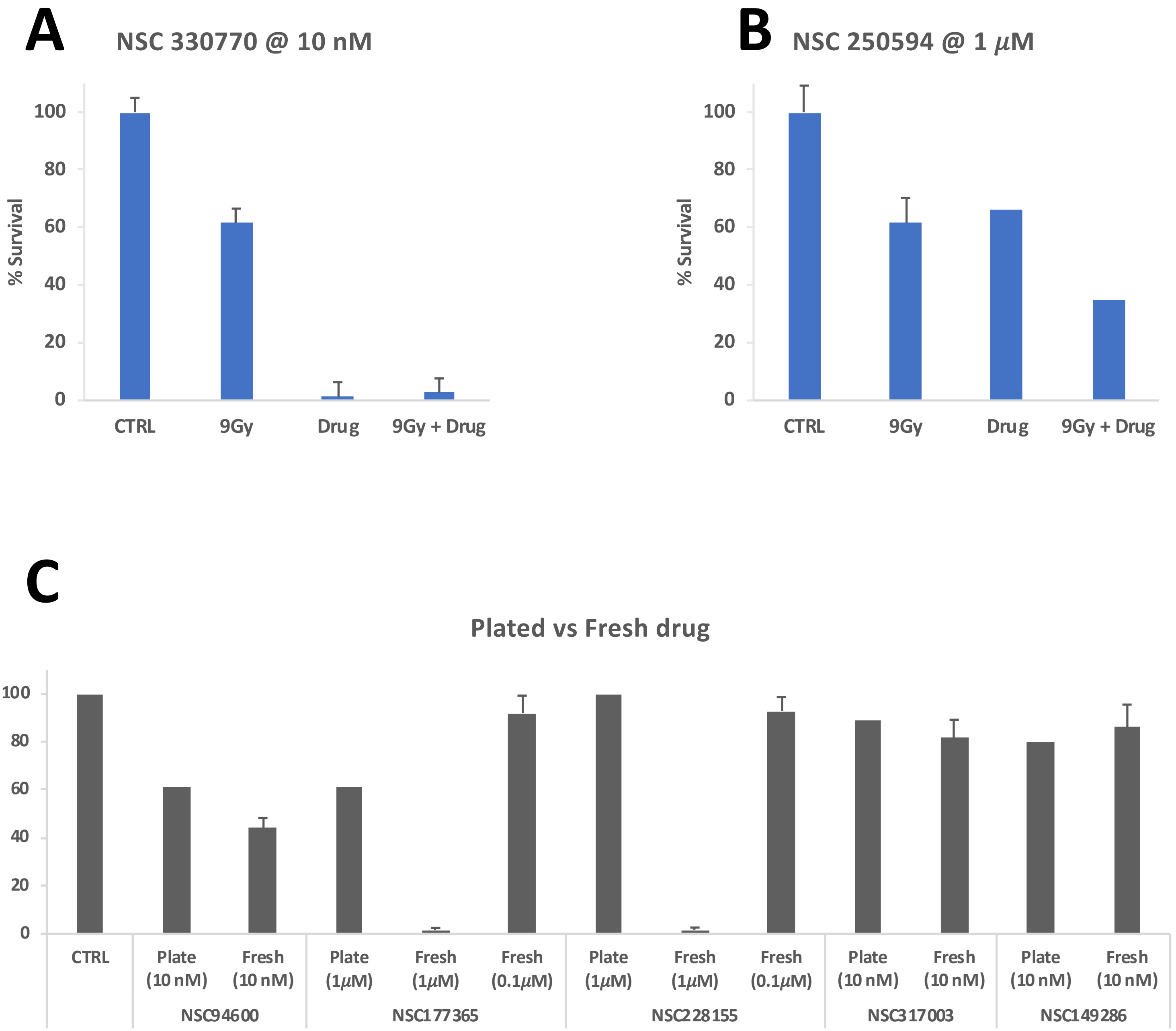


**Supplemental Figure 5**.
(A) An example of a compound that shows strong single agent activity.
(B) An example of a compound that shows additive interaction with radiation.
(C) Growth inhibition by plated and fresh compounds were compared in side-by-side testing on SW837 cells. For some (NSC94600, NSC317003 and NSC149286) the efficacy was similar between plated and fresh compounds. For others (NSC177365 and NSC228155), the efficacy was notably greater for fresh compound.

### Supplemental Figure 6.

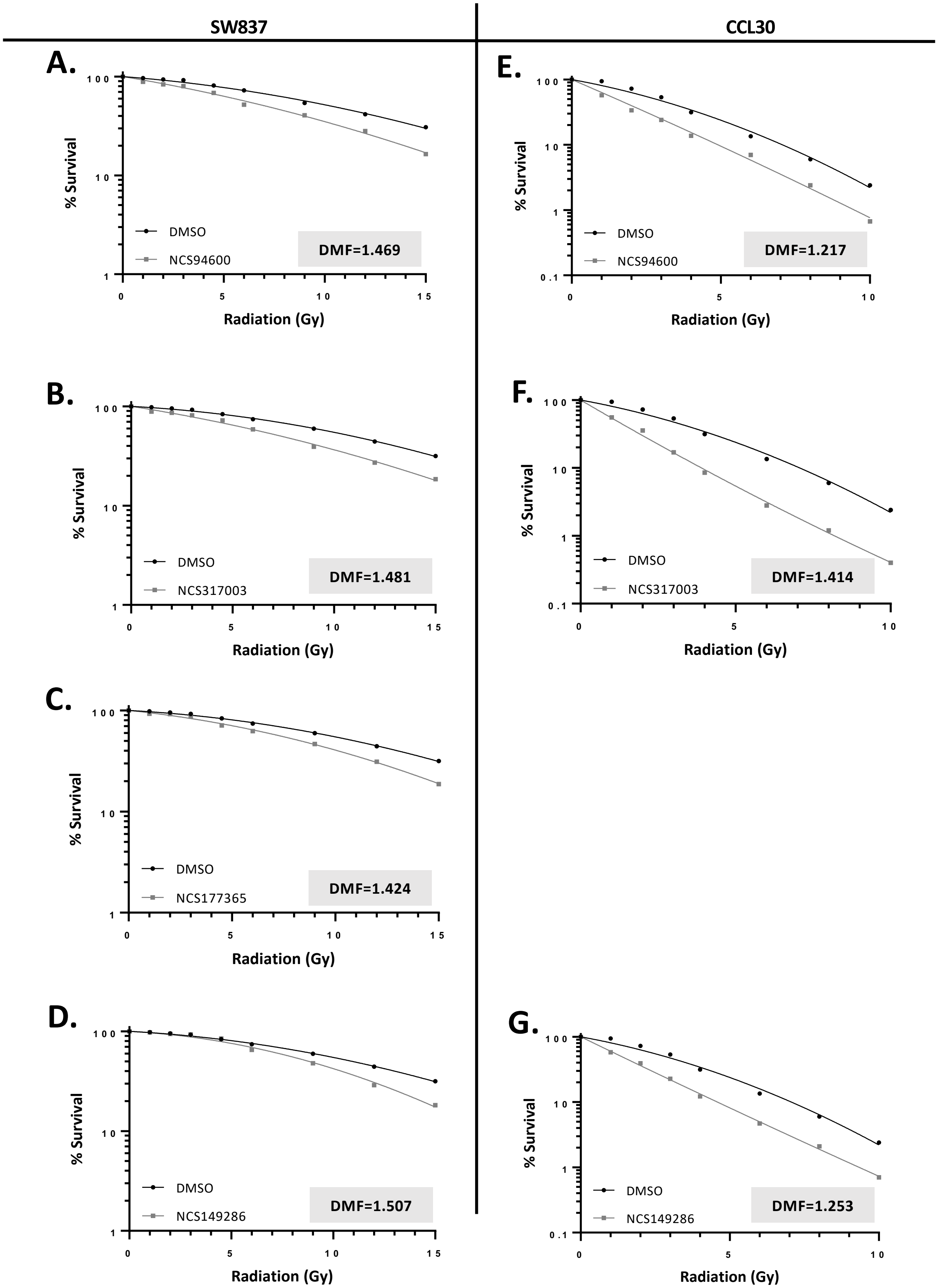


**Supplemental Figure 6**. Fresh compounds show radiation dose modification in 96-well format (as in the primary screen) on SW837 (A-D) and CCL30 (E-G) cells. Drug concentrations used with SW837 cells were 8 nM (NSC94600), 15 nM (NSC317003), 250 nM (NSC 177365) and 150 nM (NSC149286). Drug concentrations used with CCL-30 cells were 1 nM (NSC94600), 10 nM (NSC317003) and 100 nM (NSC149286). DMF was calculated as in Figure 2.

### Supplemental Figure 7

**A** **NSC317003**

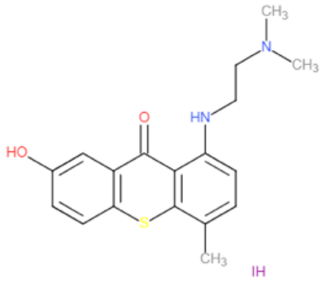


**lucanthone**

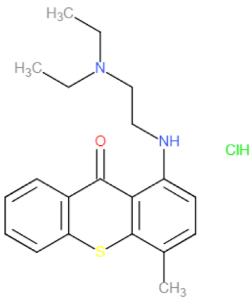


**hycanthone**

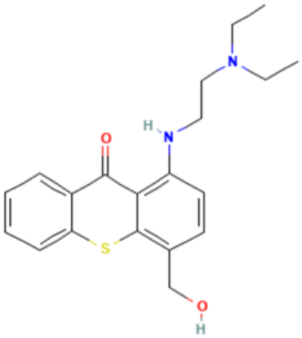


**B** **NSC149286**

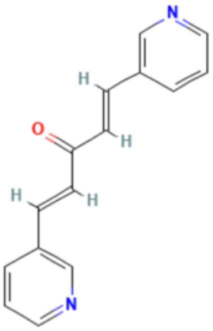


**Curcumin**

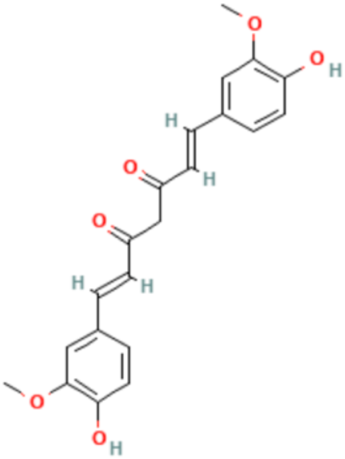


**Supplemental Figure 7**. Structures of two hits from the screen and their better-known analogs.

#### Supplemental Table 1.

**A.** Summary of the results from the secondary screen.

| Cell line | 1° hits | 2° reproducible hits | ≤ 10% survival at 10 nM | 2° additive hits | 2° modulator hits |
|---|---|---|---|---|---|
| SW837 | 41 | 39 | 6 (14.6%) | 29 (70.7%) | 4 (9.8%) |
| CCL30 | 41 | 39 | 5 (12.2%) | 31 (79.5%) | 3 (7.3%) |

**B.** Five potential radiation modulators.

| | Compound | Known or suspected activity | Extent of radio-modulation in the 2° screen | | Radiation modulator in 6-well clonogenics with fresh compound? | |
|---|---|---|---|---|---|---|
| | | | SW837 | CCL30 | SW837 | CCL30 |
| 1 | NSC94600 | Camptothecin, Topoisomerase inhibitor | >additive | >additive | YES | YES |
| 2 | NSC317003 | Structural analog of lucanthone, a DNA intercalator and topoisomerase I/II inhibitor | >additive | >additive | YES | YES |
| 3 | NSC177365 | Multiple (see text) | >additive | additive | YES | NO |
| 4 | NSC228155 | EGFR agonist | >additive | additive | NO | NO |
| 5 | NSC149286 | a curcumin analog | additive | >additive | YES | YES |

**Supplemental Table 1**.
(A) Summary of the results from the secondary screen.
(B) Five potential radiation modulators.
